# Zika virus infection induces synthesis of Digoxin in glioblastoma cells

**DOI:** 10.1101/174441

**Authors:** Estela de O. Lima, Tatiane M. Guerreiro, Carlos F. O. R. Melo, Diogo N. de Oliveira, Daisy Machado, Marcelo Lancelloti, Rodrigo R. Catharino

## Abstract

Recently, microcephaly cases have increased in Americas and have been matter of concern due to Zika virus (ZIKV) recent outbreak. Previous studies have shown that ZIKV-infected progenitor neuronal cells present morphological abnormalities and increased rates of cell death, which may be indicators of microcephaly causes. As recent studies indicate Zika virus’ tropism for brain cells, how would a glioblastoma (GBM) lineage behave under ZIKV infection, considering GBM the most common and malignant brain tumor in adults, presenting extreme chemoresistance and high morbidity and mortality rates? The current trend of using genetically engineered oncolytic pathogens as a safe way to eliminate tumors is under development, with trials already in course. Therefore, the present study evaluated the possible oncolytic effects and metabolomic alterations of Zika virus infection at human malignant M059J glioblastoma cells. Microscopic evaluation was performed using optical microscopy, which showed cytopathic effects induced by ZIKV at GBM cells. For the metabolomics study, both control and infected cell cultures were submitted to MALDI-MSI analysis. Mass spectrometry data were submitted to PLS-DA statistical analysis, and distinct biomarkers were elected for each infected groups. This study brings light to unexpectedly induced metabolic changes, as endogenous Digoxin as important biomarker for ZIKV-GBM group, associated with cytopathic effects induced by viral infection. These results evidences that genetically engineered ZIKV might be a potential new strategy for neural cancer management through the induction of endogenous digoxin synthesis in glioblastoma cells.

## INTRODUCTION

The recent outbreak of Zika virus (ZIKV) in Brazil and its outspread through the Americas has brought much concern on the effect that the infection might have on the long term ^1^. Zika virus belongs to *Flaviviridae* family and is classified as an arbovirus, a descriptive term for RNA viruses transmitted by arthropods, remarkably mosquitoes of the genus *Aedes* ^2^. Although ZIKV infection is a self-limited and asymptomatic disease in 80% of infected adults ^3^, and recent scientific evidence have proposed a strong association between mothers infected with ZIKV and newborn’s microcephaly ^1b, 4^. Currently, the absence of well-established knowledge about the exact mechanisms by which Zika virus affects human brain development is stimulating researches worldwide.

A recent literature contribution has shown that human neural progenitor cells (hNPCs) infected with ZIKV present increased rates of cell death and deregulation of cell-cycle progression, in addition to transcriptional deregulation associated with apoptotic pathways ^5^. Cell death, along with growth impairment and morphological abnormalities, was also detected by Brazilian researchers who studied Zika virus infection, Brazilian strain (ZIKV^BR^), in hNPCs ^6^. Considering the evidence that ZIKV has tropism for brain cells, induces apoptosis, and modifies cell-cycle regulation, what would Zika virus infection cause in neural cancer cells?

The 17^th^ most frequent cancer class worldwide are central nervous system tumors; out of those, glioblastomas (GBM) are the most common malignant brain tumor in adults, presenting extremely high morbidity and mortality ^7^. After diagnosis, the prognosis for GBM is generally poor, with patients’ presenting an average survival of about 11.5 months; additionally, the overall survival rate after treatment corresponds to less than 10% ^8^. Classified as grade IV gliomas, GBMs typically present increased cellular proliferation rate, invasiveness, microvascular proliferation and necrosis, all significantly more intense when compared with more lenient gliomas ^9^. The traditional treatment for GBM consists of maximal surgical resection, chemotherapy and radiotherapy. Neverthless, about 90% of surgical resections tumors recur, attesting the surgical incurability of GBMs and reduced life expectancy for these patients ^10^.

Taking into account the absence of an effective treatment for GBMs, and the ZIKV tropism for brain cells, along with its ability to induce neural cell death, the hypothesis formulated by the present contribution was that ZIKV would provoke cell death in glioblastomas through metabolic changes induced by the viral infection. Therefore, this study aimed at evaluating the metabolomic changes associated with ZIKV infection in glioblastoma cells. In addition, we proposed potential biochemical markers associated with cell death, so that future initiatives may benefit from these characteristics to interfere and improve the approaches for neural cancer treatment.

## EXPERIMENTAL SECTION

### Cell culture and viral infection

Human malignant M059J glioblastoma cells (ATCC Cat# CRL-2366, RRID:CVCL_0400), kindly provided by Professor Marcelo Lancellotti, were seeded in cover slips of a 24-well plate, cultured in RPMI medium and incubated at 37°C with 5% of CO_2_. Upon 100% of confluence, ZIKV group was submitted to viral transduction with 1x10^4^ PFU of Zika virus per well, while the control group (CT) underwent the same conditions, except for viral inoculation. Eventually, the cell culture resulted in six biological replicates for each condition studied. The viral strain obtained for this study corresponded to the Brazilian ZIKV strain (BeH823339, GenBank KU729217) isolated from a patient in 2015, at the State of Ceará, Brazil. Visual microscopic examinations were performed at 24 and 48 hours post-infection, and bright field images were collected using a Zeiss Observer A1 microscope and processed by Zeiss’ AxioVision 4 software. Finally, both groups of cells were submitted to matrix laser desorption/ionization mass spectrometry imaging (MALDI-MSI) analysis at each time point.

### Mass spectrometry analysis

At each time point of infection, 24h and 48h, cell culture samples (6 cultures for each condition, all of which analyzed in five different areas forming a total of 30 replicates for each condition) were transferred to glass slides of 24x60 mm and covered with a 10 mg/mL solution of MALDI matrix a-cyano-4-hydroxycinnamic acid (Sigma-Aldrich, St. Louis, MO) in 1:1 acetonitrile/methanol. Spectra were acquired using a MALDI LTQ-XL (Thermo Scientific, San Jose, CA) at the mass range of 400 to 1400 *m/z*, in the negative ion mode, comprising five analytical replicates per group. Tandem mass spectrometry data (MS/MS) was acquired in the same instrument, using Helium as the collision gas, with energies for collision-induced dissociation ranging from 60–140 (arbitrary units). Spectra were analyzed using XCalibur software (v. 2.4, Thermo Scientific, San Jose, CA).

### Structural elucidation

Chemical markers were elucidated via the analysis of the MS/MS fragment profile of the ions that were selected by statistical analysis. Structures were proposed with theoretical calculations modeling for molecular fragmentation with the assistance of Mass Frontier software (v. 6.0, Thermo Scientific, San Jose, CA). Structural confirmation of the cardiac glycoside Digoxin was carried out by comparison with the MS/MS profile from the analytical standard of the compound.

### Semi quantification by MSI

Chemical images of Digoxin were generated using the imaging feature of MALDI, and processed to grayscale using the ImageQuest software (Thermo Scientific, San Jose, CA). Each respective replicate from the control and ZIKV groups had their chemical image standardized and analyzed using ImageJ software (National Institutes of Health, USA—open source). A non-dimensional value was assigned to each image based on pixel intensity, so that the intensity/quantity ratio is established ^11^.

### Statistical analysis

Partial least squares discriminant analysis (PLS-DA) was used as the method of choice to check for association between groups. PLS-DA is a supervised method that uses multivariate regression techniques to extract the characteristics that may evidence this association. The selection of lipids that were characteristic for each group was carried out based on the impact that each feature had in the model, i.e. the analysis of VIP (Variable Importance in Projection) scores. As a cutoff threshold, only the chemical markers with a VIP score greater than 1.8 were analyzed. All analyses involving PLS-DA and VIP scores were carried out using the online software MetaboAnalyst 3.0 ^12^. The semi-quantitative dataset was validated with a Student’s *t*-test, and a *p*-value was significant if presented values lower than 0.05. These data were processed using the software GraphPad Prism (v.3.0, GraphPad Software, San Diego, CA).

## RESULTS

### Cytopathic effects of ZIKV in GBM cells

To identify ZIKV effects in neural cancer cells, human malignant M059J glioblastoma cells (GBM) were infected with the Brazilian strain of Zika virus. Plates were analyzed at 24 and 48 hours after infection. At the first time point evaluated (24 hours post-infection – hpi), GBM-ZIKV group (Fig. 1b) presented slight cytopathic effects, as round and swollen cells, beyond syncytium formation, compared to GBM-CT group (Fig. 1a), whilst 48hpi, GBM-ZIKV group (Fig. 1d) presented higher quantity of round and swollen cells, syncytium formation and pronounced loss of cellular integrity compared with the GBM-CT group (Fig. 1c), highlighting important cytopathic effects and cell death provoked by viral infection at the second time point of analysis.

**Figure 1.**
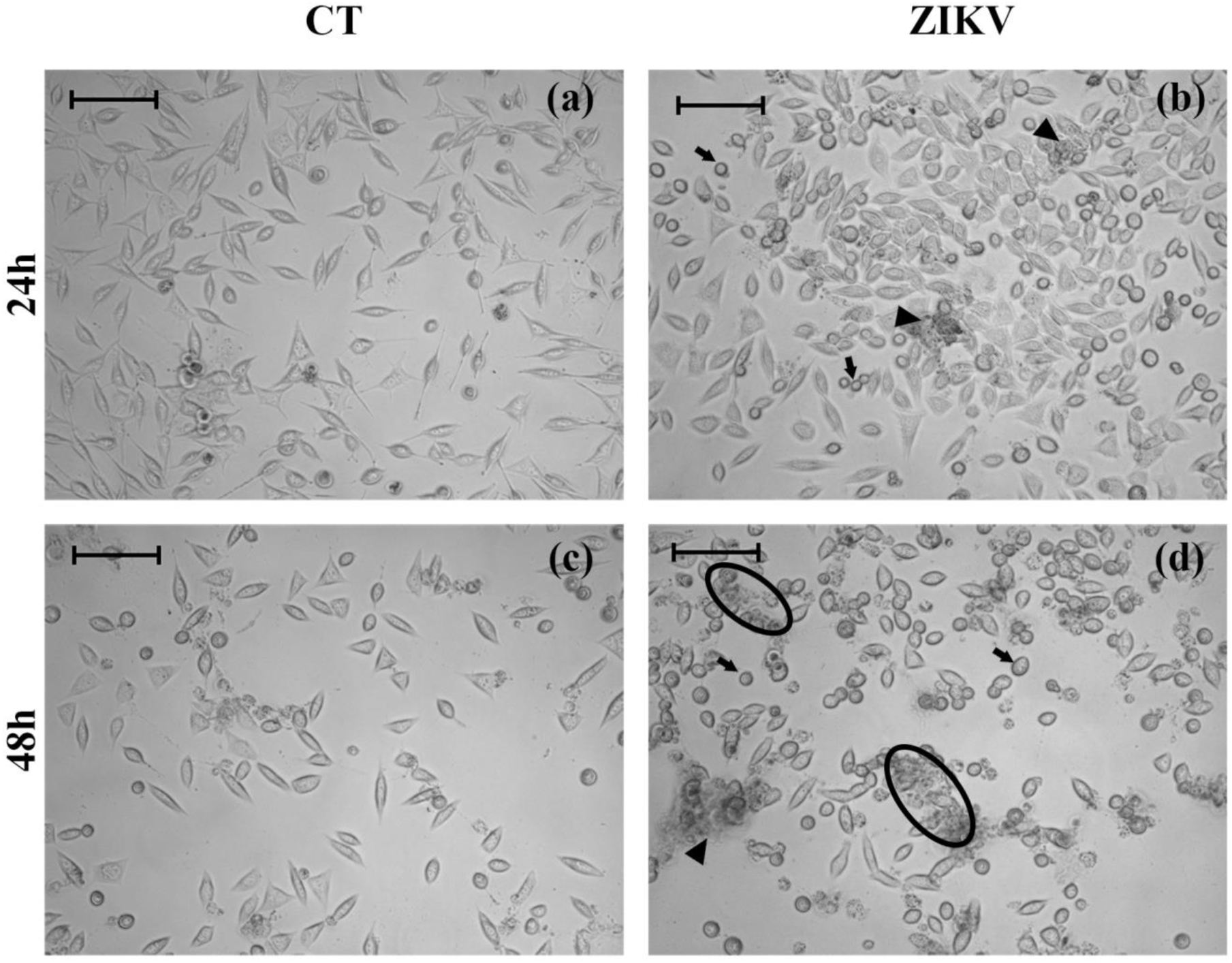
Zika virus induces cytopathic effects in glioblastoma cells.

Optical photomicrographs of glioblastoma cells from both mock (GBM-CT) and Zika virus infected (GBM-ZIKV) groups, observed at 24 and 48 hours post-infection. (a) and (b) are microscopic images made at 24 hours post infection of GBM-CT and GBM-ZIKV samples, respectively; (c) and (d) are their counterparts, 48 hours post infection. Arrows indicate round and swollen cells; arrowheads points out to syncytium formation; ellipses demonstrate areas of cytopathic effect, where there is pronounced cellular loss of integrity. Scale bars 400x, 100 μm.

### Metabolites identification through MALDI-MSI analysis

Metabolomics analytical approach was performed for both glioblastoma CT and ZIKV groups, aiming at analyzing the main metabolites involved with ZIKV infection. Mass spectrometric data (Fig. S1) were submitted to a PLS-DA, which evidenced differences between metabolite composition in ZIKV-infected cells versus uninfected control cells, for both time points of culture, as demonstrated in Fig. 2. By establishing a threshold value of 1.8 for VIP scores, it was possible to elect biochemical markers for Zika-virus infected group. For the group infected 24 hours post-infection, cells showed an interesting cardiac glycoside (CG) - Digoxin (Table 1).

**Table 1.** Lipid chemical markers elected by PLS-DA VIP scores and elucidated by MS/MS for glioblastoma cells infected with Zika virus (negative ion mode). The ion column represents the theoretical mass identified for each biomarker.

| Time of culture | Group | Class | Molecule | Precursor Ion ($m/z$) | Product Ions (MS/MS) | Molecule ID<sup>a,b</sup> |
| --- | --- | --- | --- | --- | --- | --- |
| 24 hpi | ZIKV | Cardiac glycoside | [Digoxin - H]<sup>-</sup> | 779.4 | 735, 761, 691, 752 | LMST01120023 |

Abbreviations: hpi, hours post-infection; ZIKV, Zika virus infected group.
a. LM: lipid maps ID.

**Figure 2.**
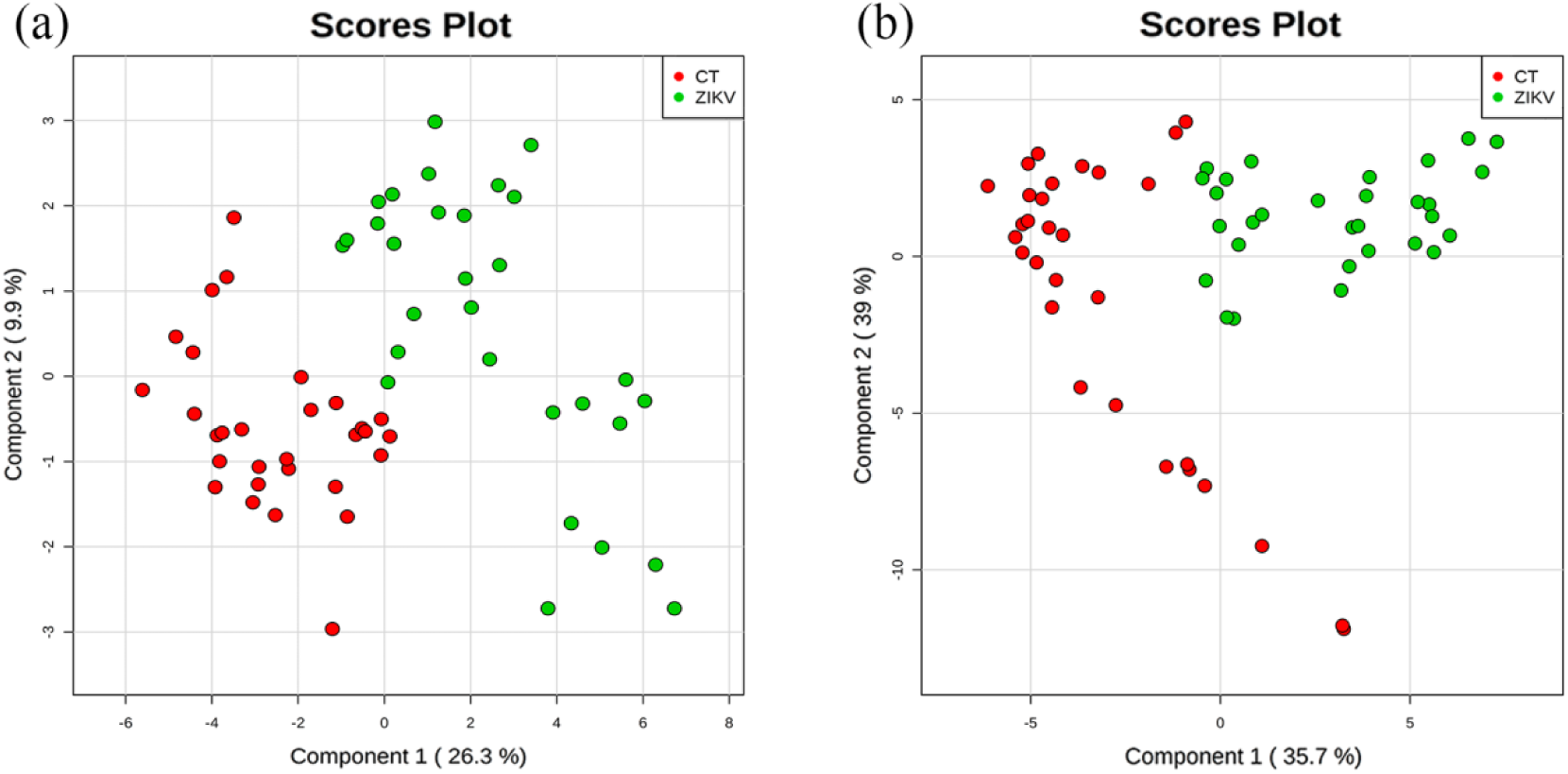
Statistical evidences of metabolomics differences between mock and Zika virus-infected groups.

Scores plot derived from PLS-DA statistical analysis between mock (CT) and Zika virus-infected (ZIKV) groups; (a) 24 hours post-infection (hpi), (b) 48 hpi. Each spot corresponds to one replicate analyzed in the cell culture, where the red spots correspond to Mock infected group and the green ones, to the ZIKV group. Each replicate corresponded to a distinct area of 10^3^x10^3^ μm in size in the cell cultures.

### Digoxin semi-quantification

Intending to corroborate the evidence that Digoxin was statistically important for the GBM-ZIKV group, the chemical image for the ion *m/z* = 779 (Fig. 3a) was semi-quantified. The statistical analysis is represented in Fig. 3b, providing enough evidence that Digoxin was significantly more expressed in Zika virus infected groups, compared to GBM-CT groups.

**Figure 3.**
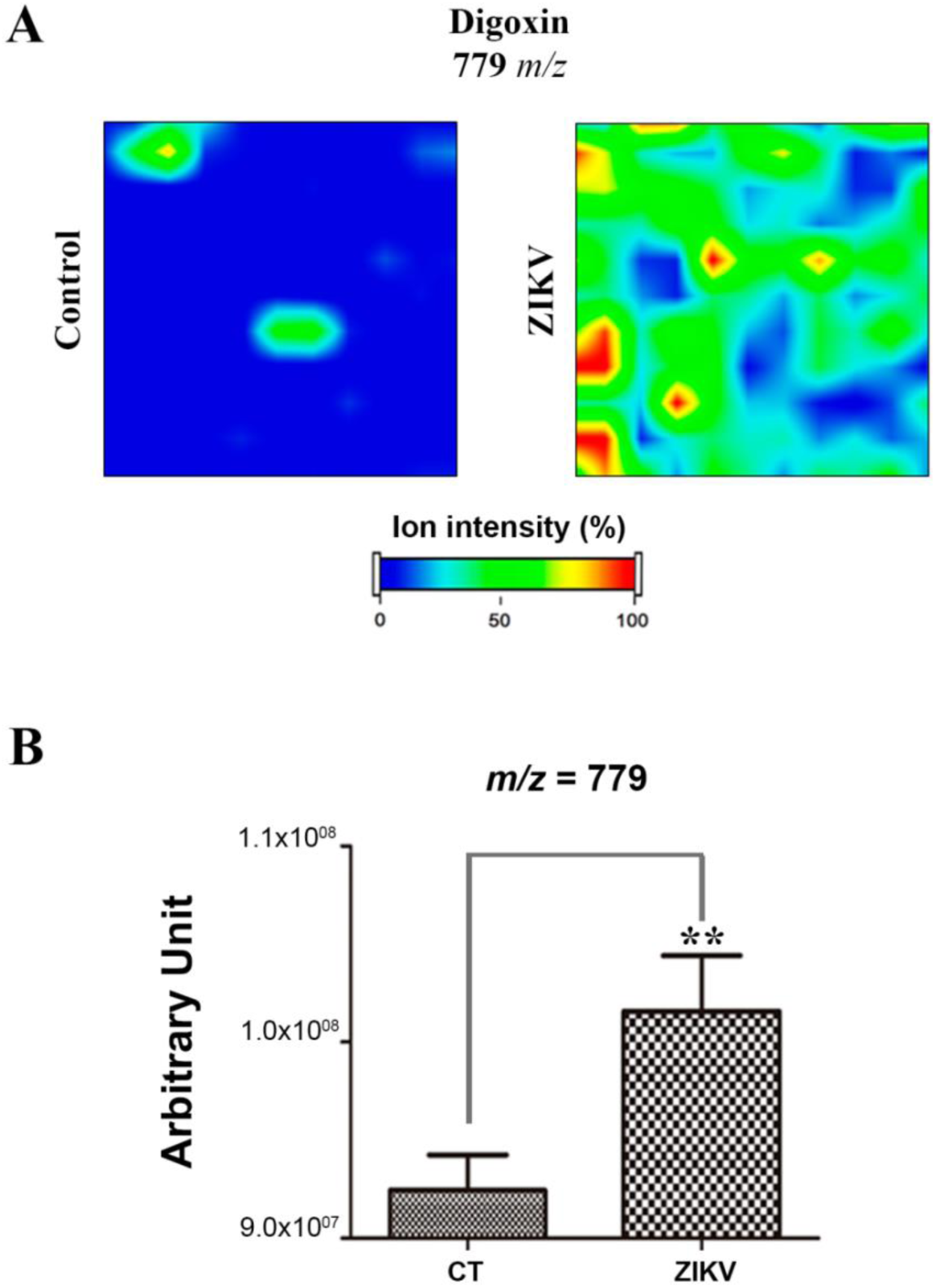
Digoxin is statistically more evident in Zika virus-infected group, 24 hpi.

(a)Digoxin (*m/z* = 779) ion image acquired from mock (control-CT) and Zika virus-infected (ZIKV) groups; (b) Semi-quantitative analysis of Digoxin observed in CT and ZIKV groups, with p = 0,002 (**).Differences on bar graphs were expressed by pixel intensities in the image for each group, as given by Oliveira *et al*., (2013). Data presented as mean ± SEM, n = 30, Student’s t-test, **p < 0.01.

## DISCUSSION

Current research points out to metabolic changes during ZIKV infection which permitted to observe an endogenous molecule that may bring light to alternative in aggressive cancer management. One of the most interesting results obtained here is that malignant glioblastoma cells are indeed susceptible to ZIKV infection, and show cytopathic effects such as altered shape, syncytium formation and loss of cellular integrity in the culture plate, all of which denote cell death, over relatively short infection times. Our findings are in line with a previous contribution that used microscopic analysis of ZIKV-infected hNPs, in which cytopathic effects were evident as early as day 2, and cell death was correlated with round and swollen shaped cells, and caspase-3 activation ^13^. Similar results were further shown for hNPCs infected with ZIKV, in which cell death was observed in association with caspase-3 activation, involved in the regulation of apoptotic pathway ^5^. This combined evidence, along with the current findings in our study, corroborate to the hypothesis that ZIKV triggers the death of neural cancer cells.

Despite the extensive research available to date, cancer management still poses a huge challenge for healthcare professionals worldwide. Therefore, several members of the biomedical community have been engaged in better understanding the biochemical basis and the physiology of cancer, intending to improve the performance of cancer treatment. For this purpose, several studies are emerging and bringing light to novel and unusual data about new alternatives for cancer treatment. Within this scope, a class of molecules has been closely monitored based in clinical and scientific evidences: cardiac glycosides (CGs) ^14^.

The most interesting and intriguing molecule identified in GBM-ZIKV group at the early stage of infection (24 hours post-infection) belongs to cardiac glycosides group, more specifically the cardenolides subgroup ^15^, and consists of Digoxin. Cardiac glycosides are sterol lipids, well known as secondary metabolites in plants, however, there are few studies that describe these molecules as endogenous compounds in mammals ^16^. By the end of the past century, scientific studies, including mass spectrometric analyses showed that cardiac glycosides could be identified in human tissues and fluids, such as adrenal glands, kidneys, brain, plasma and urine ^17^. Endogenous *de novo* biosynthesis of CGs was firstly reported by Qazzaz and colleagues (2004), attesting the previous evidences cited above ^17a, 18^. Furthermore, previous studies have demonstrated that patients with breast cancer, under cardiac glycosides treatment present reduced recurrence and mortality rates compared with the untreated group ^19^. *In vitro* studies have shown that innumerous cancer cell lines, like breast, melanoma and neuroblastoma cells present reduced multiplication rate and increased cell death after treatment with CGs ^20^. In addition, cardiac glycosides like Digoxin were also evaluated regarding GBM cells response, and the treatment induced proliferation impairment and apoptotic phenotype ^21^. Considering the existence of several *in vivo* and *in vitro* data showing that CGs induces cancer cell death, it is plausible to admit that ZIKV infection brought on endogenous synthesis of Digoxin in glioblastoma cells. This contrasts directly with the group with mock-infected cells, where the presence of this cardiac glycoside was not observed. Thus, we propose that this phenomenon is probably one of the triggers for neuronal cell death induced by Zika virus.

Digoxin is a well-known potent inhibitor of the Na^+^-K^+^-ATPase ion pump, which is responsible for the establishment and maintenance of the electrochemical gradient across plasma membrane ^22^. Taking into account that neuronal signaling depends on generation and maintenance of membrane potential, Digoxin may disrupt ion homeostasis and disturb neuronal excitability, both for healthy and cancer neuronal cells. Furthermore, the sodium-potassium pump inhibitory effect leads to intracellular increase in sodium and calcium levels, which mediates several signaling pathways, including proliferation, differentiation and apoptosis ^22b, 23^. Although Na^+^-K^+^-ATPase is an enzyme present in every mammalian cell, in this case, where synthesis of digoxin is induced by Zika virus infection, the harmful effects of cardenolides would be restricted to infected cells, what would protect the non-infected ones.

Beyond the effects cited above, some studies have shown possible mechanisms of action for cardiac gangliosides in cancer cells, suggesting they might interfere in different signaling pathways, what initiates with their interaction with Na^+^-K^+^-ATPase ion pump. This first step triggers changes at different signaling cascades via distinct proteins, as Src (Proto-oncogene tyrosine kinase), Phosphoinositide 3-kinase (PI3K), Phospolipase C and Ras pathways, which together might result in tumor cell death through apoptosis or autophagy mechanisms ^24^. Some studies have also shown that digoxin induces apoptosis via activation of Caspase-3, which ends with generation of reactive oxygen species (ROS) and cell death ^25^. All these signaling cascades together will affect DNA translation and consequently alter synthesis of proteins which are involved with growth, survive and cell cycle control ^26^. Digoxin, for instance, showed reduction of hypoxia-inducible factor 1 (HIF-1α) gene expression and protein synthesis at cancer cells, which impairs the role of HIF-1α as an important transcription factor for cancer angiogenesis. Although digoxin inhibition of HIF-1α occurs, HIF-1α expression is not affected when cells are treated with a vector containing only coding sequences, generating mRNA without untranslated regions (UTRs). That means that digoxin might function as an mRNA translational regulator at UTRs from HIF-1a mRNA generated ^27^. In case of cancers, negative interference with vascular support for GBM cells is of great value for cancer management, and digoxin induced by Zika virus might be an alternative to weaken tumor cells and reduce glioblastoma growth. Although the exact mechanism of action of cardiac glycosides still needs to be elucidated, Digoxin seems to be a robust candidate for neural cancer management.

Although Digoxin was not elected as a significant biomarker for GBM-ZIKV group in the later stage of infection (48 hours post-infection), it indicates that Digoxin was important at the first hours after ZIKV infection. It is likely Digoxin, in the first 24 hours of infection, was synthesized and induced intracellular changes that triggered cell death pathways after 48 hours. The reduction of living cells after 48 hpi impairs digoxin election as a statistical biomarker. Furthermore, the biochemical changes caused by Digoxin might have overlapped its own expression and reduced its significance at statistical analysis.

Therefore, the present study has shown that malignant glioblastoma cells were susceptible to ZIKV infection, which was verified through cytopathic effects over infection time. Therefore, microscopic observations combined with the metabolomics data revealed in the present study suggest that Digoxin, synthesized after stimulation by Zika virus, may be responsible for glioblastoma cells death. Based on these results, remains the question whether a genetically engineered Zika virus might be an alternative for cancer treatment.

Since Zika virus presents tropism for neural progenitor cells, originated from either fetal or adult brains ^13, 28^, the consequences of the infection in adults are not as severe as in fetuses ^28^, as fetuses present a higher number of neural stem cells compared to adults ^29^. This vulnerability of neural precursors to Zika virus has recently been confirmed by Chavali *et al*., 2017. This study demonstrated that cells with higher expression of protein Musashi-1 (MSI-1), like glioblastoma and neuronal precursor cells, presented high viral replication rates. The presence of MSI-1 was essential for Zika virus replication, once MSI-1 was responsible for promoting viral RNA translation in stem cells ^30^. Thus, researchers concluded that Zika virus might present tropism for immature neuronal cells because MSI-1 is highly expressed on it, but absent at totally differentiated neuronal cells, what explains the deleterious effects of ZIKV infection at fetuses’ brains, differently of adult’s brains. Considering that glioblastomas might be originated from cancerous neural stem cells (immature cells) ^31^, it is plausible to consider them as important Zika virus targets.

Therefore, through the sensitivity of glioblastoma cells to Zika virus infection demonstrated in the present study, and the viral selectivity for progenitor cells, we propose that the synthesis of Digoxin induced by Zika virus might be an innovative alternative for neural cancer management. In a practical example, members from the Food and Drug Administration – FDA and European Medicines Agency (EMA) have already approved the use of genetically engineered virus, called Talimogene laherparepvec (T-VEC) to treat advanced melanoma ^32^. Clinical trials with different oncolytic viruses are already in course, as in the case of avian adenovirus, myxoma virus, feline panleukopenia virus, polio virus, among others ^33^. An interesting case was observed during a clinical trial with an engineered measles virus, where a myeloma patient entered on complete remission after the treatment 34. One of the reasons for these trials is that many viruses preferentially infect cancer cells, and if so, healthy cells could be safe while viral infection eliminates the tumor ^32b^. Therefore, the use of oncolytic viruses is a promising strategy, and further studies must be taken forward to verify the link between synthesis of Digoxin and Zika virus infection. ZIKV may be, therefore, ultimately genetically engineered and be a strong alternative to glioblastoma management, in an effort to provide better quality of life and survival rates for these patients.

In addition to the possibility of using ZIKV for neural cancer management, an alternative to explore the results obtained from this research is to evaluate the potential use of Digoxin for Zika virus diagnosis. Current ZIKV diagnostic tools are based on the detection of antibodies (immunoassays) and/or viral components, whether viral proteins or RNA. The routine diagnostic method for ZIKV infection is RT-PCR, which might be complemented with MAC-ELISA assay ^35^. However, the low levels of viremia after one week of the illness onset and the cross-reactivity of flavivirus antibodies makes difficult to obtain the correct diagnosis ^36^. Taking into account the need for highest specificity and sensitivity for ZIKV diagnosis, Digoxin might be a potential biomarker for that. Further studies may be conducted to verify the presence of this chemical marker in biological samples from infected patients, in addition to evaluate the specificity of cardiac glycosides for ZIKV infection. Once Digoxin is validated as a ZIKV biomarker, it is possible to ultimately apply the metabolomics strategy for the diagnosis of Zika virus infection.

## ACKNOWLEDGMENTS

Innovare Laboratory would like to thank Professor José Butori Lopes de Faria and Diego Andreazzi Duarte for providing the infrastructure of the microscopy lab of the *Laboratório de Microdiabetes from the Clínica Médica (FCM)*, essential for cell imaging. We also acknowledge São Paulo Research Foundation (FAPESP, Process Nos. 11/50400-0 and 15/06809-1 for RRC, 14/00302-0 for CZE. We also thank Coordination for the Improvement of Higher Level Personnel (CAPES) for the fellowships from CFORM (PROEX: 1645986), EOL (PNPD: 1578388) and TMG(PROEX: 1489740). DNO acknowledges the Plano Nacional de Enfrentamento ao Aedes aegypti e à Microcefalia [*Brazilian Plan for Fighting Aedes aegypti and Microcephaly*] for the fellowship under process No. 88887.137889/2017-00. All researchers involved declare no conflicts of interest of any nature.

## AUTHOR INFORMATION NOTES

Corresponding author: Phone: 55 19 35219138

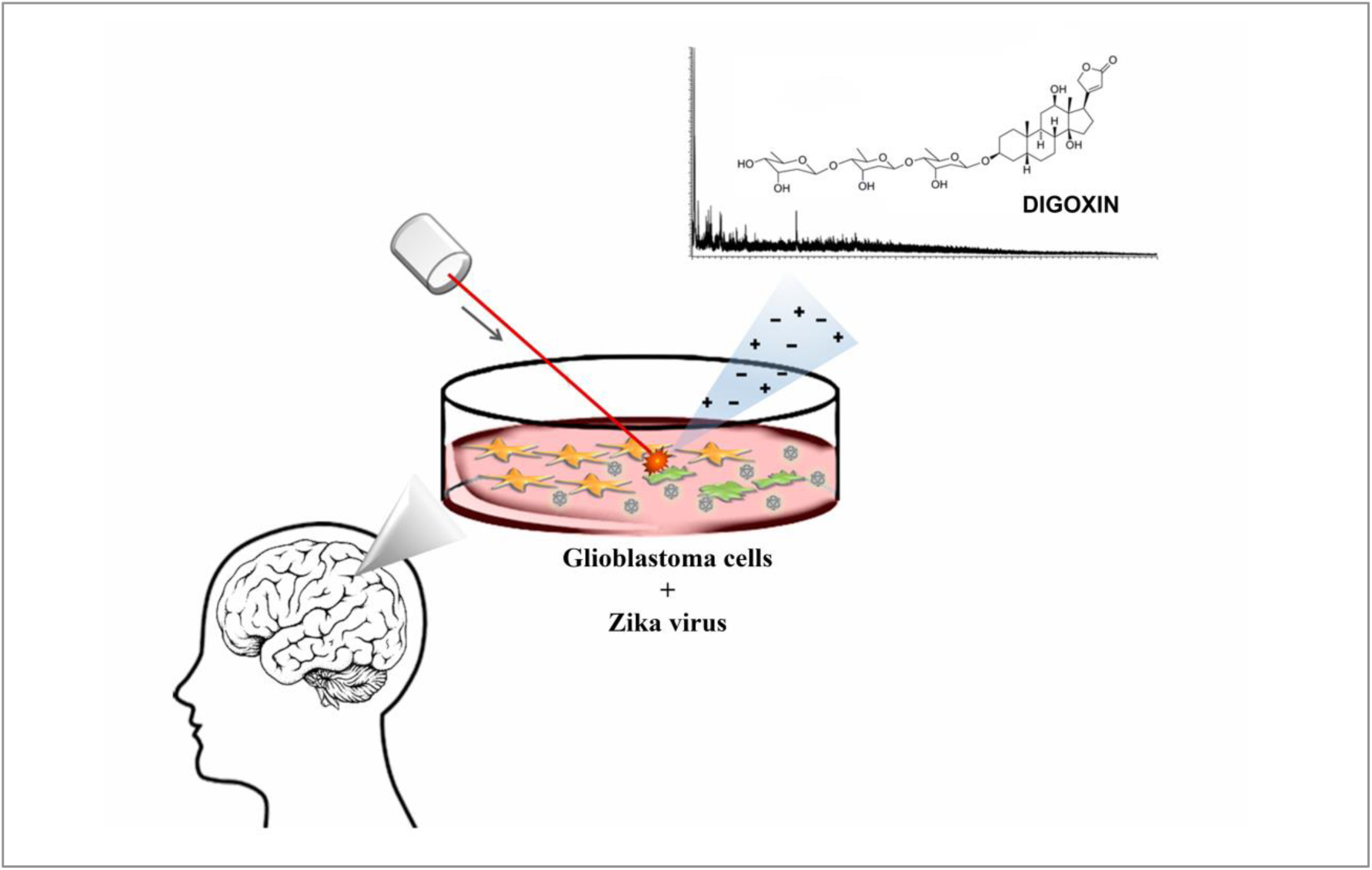

